# Local alkalinity enhancement using artificial substrates increases survivorship of early-stage coral recruits

**DOI:** 10.1101/2025.01.07.631763

**Authors:** Melissa Ruszczyk, Skylar Rodriguez, Montale Tuen, Kylee Rux, Santhan Chandragiri, Maren Stickley, Brian K. Haus, Andrew C. Baker, Margaret W. Miller, Prannoy Suraneni, Chris Langdon, Vivek N. Prakash

## Abstract

Efforts to restore coral reefs using sexually derived coral recruits are often hindered by their low survivorship and growth, hence scalable interventions to improve these parameters are urgently needed. Here, we developed novel settlement substrates that modify the local chemical and hydrodynamic environments to provide local alkalinity enhancement (AE) within the laminar boundary layer to aid in coral restoration. Cement tiles with four different chemistries and two different surface topographies were tested in a novel flume system to quantify their ability to change local pH under reef-like conditions and their effect on larval settlement, survivorship, and growth of the endangered Caribbean coral, *Orbicella faveolata*. Chemistry had a minimal effect on the initial settlement of coral larvae, and textured tiles were preferred over smooth tiles. However, substrates that created a more alkaline local environment increased post-settlement survivorship. The increased survivorship of *O. faveolata* recruits on AE tiles was not due to increased growth on AE tiles, although trends in growth were dependent on chemistry and topography of tiles. Our results indicate that mixing sodium bicarbonate or sodium carbonate into cement used to fabricate artificial reef structures could be an effective means to enhance the development of coral cover.

**Significance Statement:** Reef-building coral populations in South Florida and the Caribbean are nearing crisis, with low survivorship rates of new recruits being identified as a critical bottleneck limiting their recovery. Since ocean acidification is known to stress corals and hinder their growth, we hypothesized that enhancing substrate alkalinity could help mitigate ocean acidification effects by boosting coral growth and increasing survivorship during their critical early stages. This hypothesis was experimentally tested by adding sodium bicarbonate or sodium carbonate to settlement substrates and studying coral settlement preference, and coral spat growth and survival. Our results showed significantly higher survivorship on alkalinity enhanced substrates, indicating that this could be a promising restoration intervention.

## Introduction

Coastal cities and communities are susceptible to the impacts of waves, flooding, storm surge, and sea-level rise. In response to these threats, coastal jurisdictions typically invest in engineered shoreline defenses such as breakwaters and seawalls that are expensive to build and maintain. To date, built infrastructure, including sea walls, levees, culverts, bulkheads, and other hardened structures, have dominated thinking about coastal protection (1). However, there is increasing interest in using natural habitats, including wetlands, dunes, barrier islands, sea grasses, coral and oyster reefs, and mangroves to reduce the risk of coastal flooding and erosion and provide other social and economic benefits (2). Many of the co-benefits associated with natural infrastructure are precisely what make coastal areas so valuable and what draws people to live and work in these vulnerable regions. The coastal ecosystems that enhance resilience by providing protective services also contribute raw goods and materials, plant and animal habitat, water and air quality regulation, carbon sequestration, nutrient cycling, and opportunities for tourism, recreation, and education (3).

There is growing global interest in combining the use of built and natural infrastructure to help coastal communities become more resilient to extreme events and reduce the risk of coastal flooding through the creation of hybrid coastal resilience structures. One such hybrid approach is the deployment of an artificial reef base structure consisting of a submerged breakwater onto which reef-building corals are actively restored. These hybrid reefs which incorporate an overlay of living corals could offer the immediate benefits of wave attenuation while providing ecosystem benefits and promoting ecosystem resilience (4). Coral reefs are naturally formed, low-crested breakwaters that aid in wave energy dissipation and flood reduction and represent biodiversity hot spots that provide valuable ecological and economic benefits (5, 6). Healthy coral reefs can attenuate wave energy by up to 97%, providing tremendous natural protection from coastal hazards (7). Considering this, hybrid reefs might best be thought of as opportunities to offer communities the immediate benefits of shoreline protection and the long-term benefits of restoring coral reefs, and important economic benefits and ecological services. In fulfillment of this vision, hybrid coral reef projects would become testbeds for testing new methods of coral restoration and increasing coral resilience to climate change.

While the concept of hybrid coral reefs sounds attractive, natural coral reefs around the world are diminishing rapidly. According to the latest status of the world’s coral reef report, between 2009 and 2018, there was a progressive loss of about 14% coral cover from the world’s coral reefs (8). This was primarily due to recurring large-scale coral bleaching events, combined with other local pressures such as coastal development, land-based and marine pollution, unsustainable fishing, disease, and tropical storms.

The global decline in coral populations creates a snowballing problem of decreased larval supply and decreased genetic diversity, inhibiting recovery following disturbance. Larval availability, settlement success, and post-settlement survivorship and growth have long been recognized as key factors driving the recovery of coral reefs following disturbances (9). Studies have shown that settlement rates of corals in the Caribbean are much lower than on reefs in the Pacific and Indian Oceans (10). This could be related to generally much lower coral cover on Caribbean reefs and hence a smaller supply of larvae (11). Moreover, slower post-settlement growth is an important contributing factor because low growth rates mean that early-stage recruits remain longer in the very small size classes that are most vulnerable to predation and competitive exclusion by macro-algae.

Ocean acidification is known to stress corals, causing them to grow more slowly or divert energy from other essential life processes to maintain a constant growth rate. This led us to hypothesize that alkalinity enhancement (AE) might be a way of relieving physiological stress of the coral spat. Adding alkaline compounds to the environment has been shown to increase calcification of adult corals (12-14). However, these studies involved adding chemicals to the seawater around corals. While the additions had the desired effect, this approach is not readily sustainable long-term because of the large amounts of chemicals that would need to be added given the volume of seawater that flows over a coral reef every day. Alternatively, compounds increasing alkalinity can be added to substrate where they leach into the water column, providing an alkalinity-enhanced boundary layer for coral spat (**Figure 1a**). We designed AE tiles made with four different chemistries (Portland limestone cement with no additions, 1% sodium bicarbonate by weight, 1% sodium carbonate by weight, and 2% sodium carbonate by weight, referred to as tile ‘chemistries’) and two different surface topographies (a flat tile, and a textured tile with concave divots) as substrates. These AE tiles may increase local alkalinity in the fluid above both by diffusion and advection inside a boundary layer in the laminar flows generated using a novel flume **(Figure 1b-d)** (15). Carbonate and bicarbonate ions leaching out of the tiles may be sufficient to stimulate early growth of the coral spat through either direct contact of the coral on the substrate or while the coral is small enough to occupy the laminar boundary layer (**Figure 1a**). This AE method via substrates could be less resource-intensive than bulk additions by mixing small amounts of sodium carbonate/bicarbonate into the cement used to fabricate artificial or hybrid reefs, or by the production of tiles using these chemistries for the land-based production and grow-out of coral spat.

**Figure 1.**
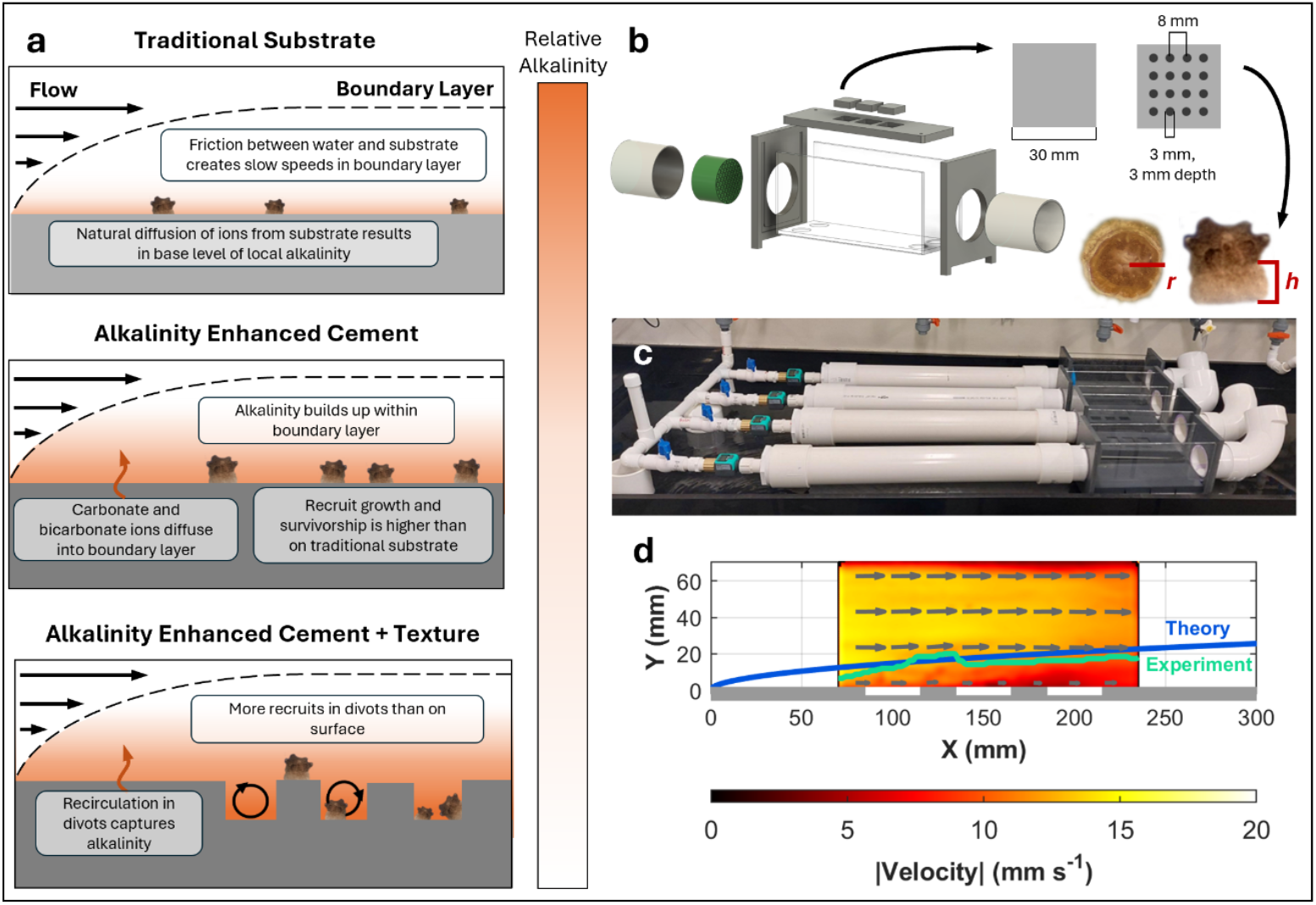
Local alkalinity enhancement concept and experiments. (a) Top panel: Coral recruits (< 1 cm in height) grow within the laminar boundary layer of a substrate, where the local flow velocity is slower than the bulk flow above. Slower velocities inside the laminar boundary layer allow for ions that diffuse from the substrate to stay near the surface for longer times. Middle panel: Alkalinity enhanced substrates leach/release carbonate and/or bicarbonate ions into the water column, resulting in a higher alkalinity environment that corals can take advantage of, resulting in more growth and greater survivorship. Bottom panel: By modifying the topography of the substrate, it is possible to create a dynamic landscape with regions of even greater local alkalinity which may further enhance coral growth and survivorship. (b) Experimental viewing chamber design and flume components: Each chamber holds three cement tiles that are either flat or textured (with surface divots). *Orbicella faveolata* recruits grow on the surface of the cement tiles. Their radius, *r*, and skeletal height, *h*, are used to determine growth measurements. (c) Four flow-through flumes connect to one inlet line allowing for multiple experimental flumes. (d) Quantification of fluid flow in the flume viewing chamber where the tiles sit. The experimentally measured laminar boundary layer (green) and the theoretical boundary layer (blue) are included.

## Results

### Tile Verification: Diffusion and Advection

Adding 1-2% sodium bicarbonate or sodium carbonate by weight to cement tiles was sufficient to elevate pH by 0.2 – 0.4 pH units under stagnant flow conditions (*P* < 0.0001) and 0.1 pH units within the laminar boundary layer generated in our flumes (15). The laminar flows in the flumes were comparable to that encountered on a coral reef (*P* < 0.0001, **Figure 2a-c**). Within the laminar boundary layer, all AE tiles significantly increased pH; however, the effect of 2% carbonate tiles under advective flow was greatly diminished compared to the diffusive (no flow) setting (**Figure 2c, Supplementary Table S1**). The smaller increase of pH in the advection setting occurred because the carbonate and bicarbonate leaching out of the cement was continually diluted by incoming seawater.

**Figure 2.**
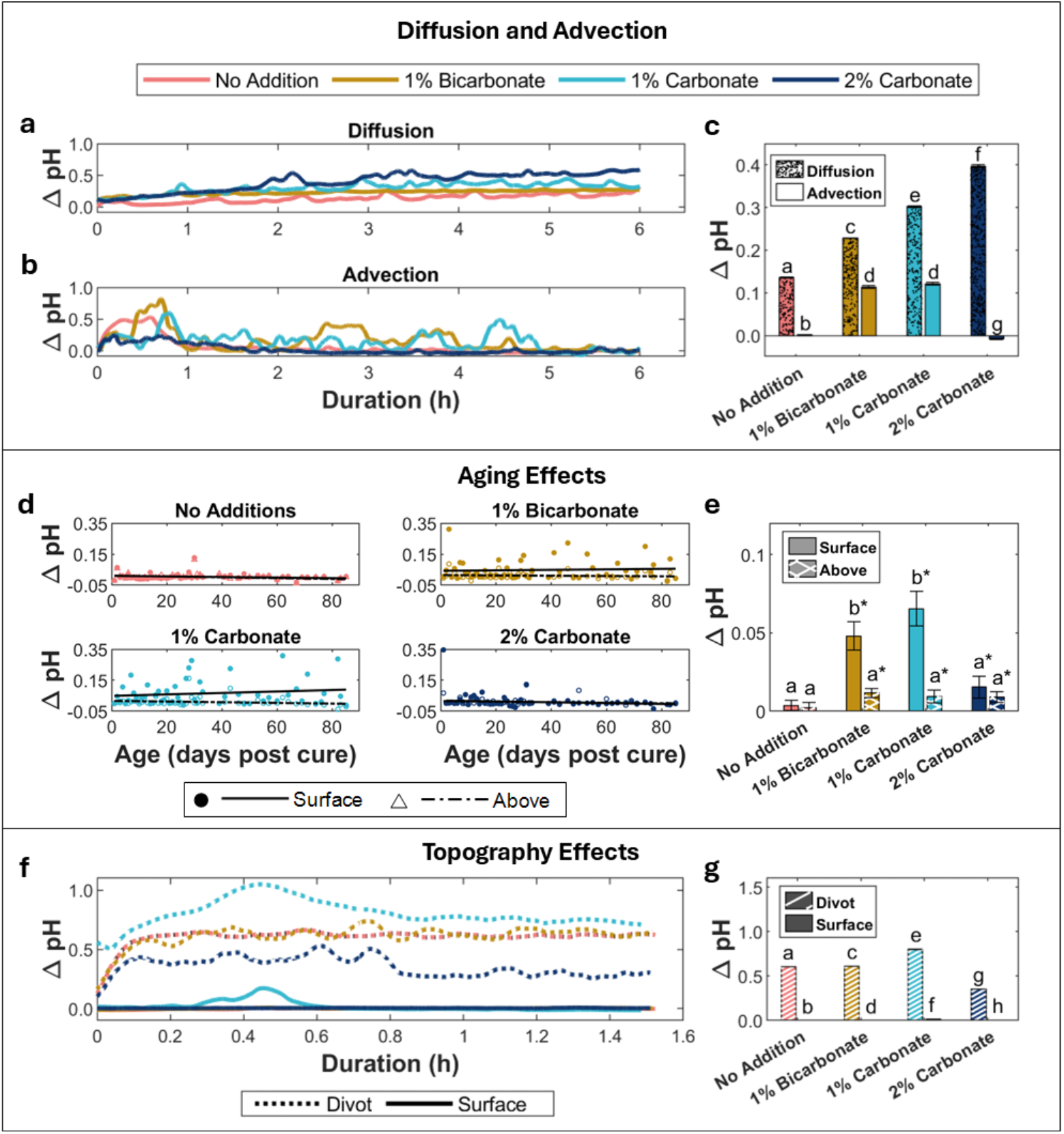
Physicochemical characteristics of AE tiles. (a) The first 6 h of a 24 h diffusion time series for flat tiles. (b) The first 6 h of a 24 h advection time series for flat tiles. (c) The average *ΔpH* from hour 1-6, after the tile reaches equilibrium with the environment. (d) Linear regressions fitted to *ΔpH* for tiles with no additions, 1% bicarbonate additions, 1% carbonate additions, and 2% carbonate additions, at the surface and 3.26 ± 0.09 mm above the surface of the tile to capture changes on the surface of the tile and within the laminar boundary layer in flow conditions. (e) Average *ΔpH* at and above the boundary layer of the four different chemistries throughout the 84-day aging experiment. Asterisks indicate average value greater than 0. (f) The first 90 min of advective time series on textured tiles measured at the surface of the tile and within a divot on a textured tile. (g) The average *ΔpH* over the duration of the advective time series for textured tiles. Throughout, different letters indicate statistically different data.

### Longevity of Tiles

Linear regressions were fitted to the change in pH for the different chemistries versus tile age (1-84 days post cure, or up to 12 weeks post cure) for measurements at the surface of the tile and 3.26 ± 0.09 mm above the surface of the tile to capture long-term changes above the surface of the tile and within the laminar boundary layer (**Figure 2d**). There was one outlier in the entire dataset noted for 2% carbonate additions that occurred on day 1 and was excluded from the analysis (but included in **Figure 2d** for reference). Three out of the eight total regressions were significant, indicating the declining effectiveness of tiles with no additions (*P*_*Surface*_ = 0.01), and 2% carbonate additions (*P*_*Surface*_ < 0.01, *P*_*Above*_ = 0.03) to increase pH over the 12-week aging experiment, while 1% bicarbonate addition and 1% carbonate additions did not change with respect to time (**Figure 2d, Supplementary Table S2**).

Data points were grouped by chemistry and measurement location over the duration of the experiment to identify broad trends. At the surface of the tile, 1% bicarbonate additions and 1% carbonate additions resulted in the greatest increases in pH (**Supplementary Table S3**). Additionally, Students’ t-tests were run to see if the lower-performing tiles (i.e., no additions or 2% carbonate additions) still increased pH compared to the baseline pH with no tile present. All tiles, except for those with no additions, had an average pH higher than baseline at the surface of the tile and within the laminar boundary layer (**Supplementary Table S3**) (**Figure 2e**). Overall, these aging tests showed that the AE tiles could elevate pH within the fluid boundary later by 0.05-0.06 units for at least 12 weeks.

### Chemistry and Topography

The tile topographies in this experiment included a flat surface (i.e., a “flat” tile), and a surface textured with divots (i.e., a “textured” tile). There were minimal differences in the flow above the tiles between these two surface topographies with regards to velocity and vorticity, indicating that there is no breakdown of the laminar boundary layer resulting from the tile divots (**Figure S5**). This was because the smooth, laminar flow within the flumes did not generate recirculation regions.

Even though there was no difference in the hydrodynamic environment between the two topographies, the chemical landscape around the tiles differed. A full factorial ANOVA comparing the change in pH with respect to the four different chemistries and two locations within textured tiles (i.e., in a divot or on the surface of the tile) indicated significant differences in ΔpH with respect to chemistry (*P* < 0.0001) and location (*P* < 0.0001), with ΔpH being significantly higher in the divots than on the surface of a textured tiles for all chemistries (**Figure 2f-g**). Because there was minimal flow into the divots resulting in minimal advection, the pH in the divots was significantly higher than at the surface. Hence, the divots provide a region sheltered from flow (*u* ≈ 0 m s^-1^).

### Settlement

To identify any settlement preference, the number of *O. faveolata* settlers per tile was analyzed on day 0 using an ANOVA, with chemistry and topography as factors. There was no significant difference in settlement based on chemistry (*P* = 0.47), however, textured tiles had higher settlement numbers than flat tiles (*P* < 0.001, **Figure 3a**). On textured tiles, where corals had the option of settling on the surface or in a divot, corals preferred settling in divots (*P* < 0.01, **Figure 3b**).

**Figure 3:**
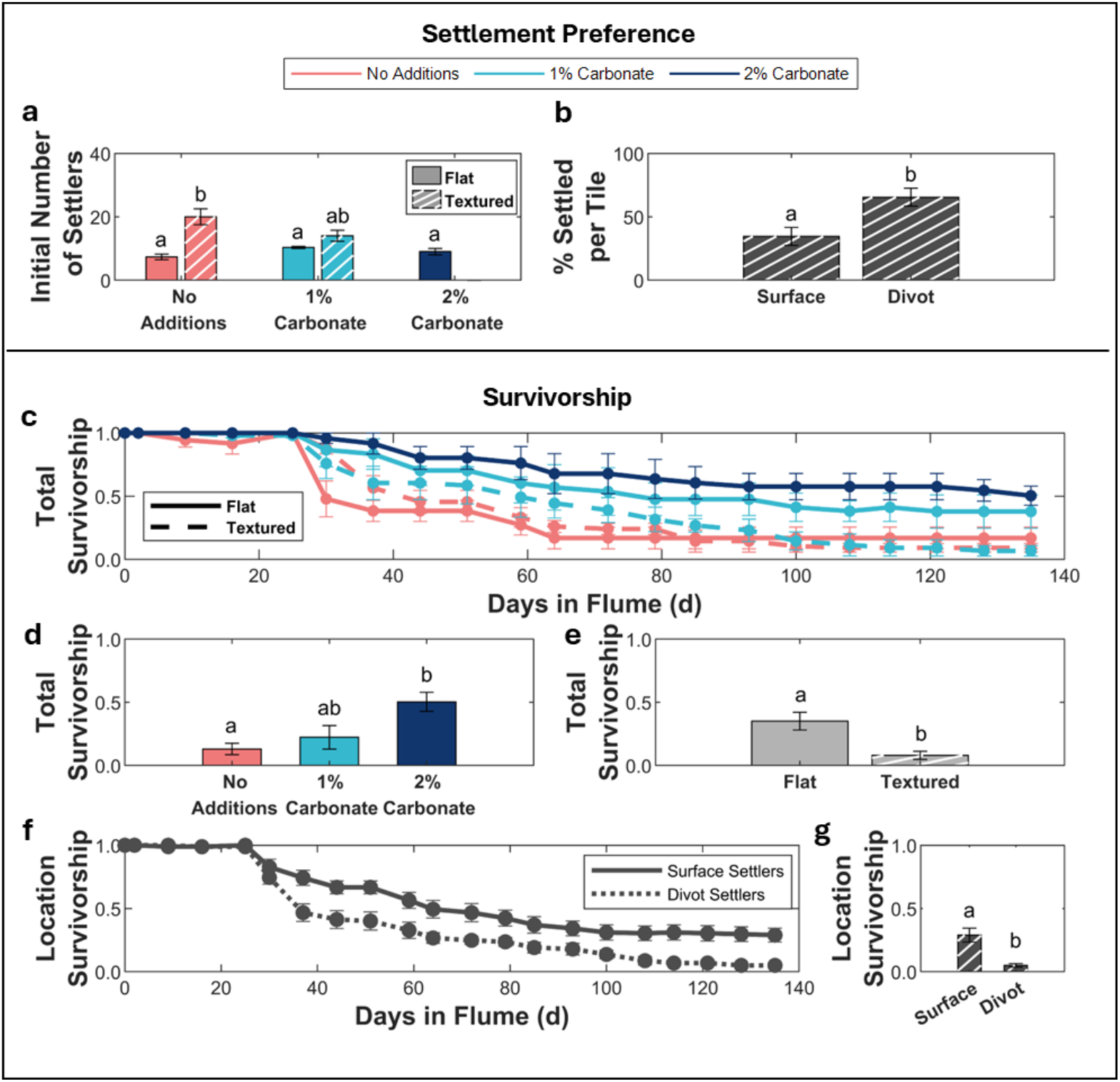
*Orbicella faveolata* settlement and survivorship. (a) The initial number of *O. faveolata* settlers on all five tile types tested. (b) The percentage of settlers based on location within the textured tiles. (c) Survivorship curves for the three different chemistries and the two different surface topographies. (d) Average total tile survivorship across all tiles (n = 15), regardless of settlement location, at the end of data collection with respect to chemistry. (e) Average total survivorship across all tiles (n = 15), regardless of settlement location, at the end of data collection with respect to topography. (f) Location survivorship curves for surface and divot settlers on textured tiles only (n = 6). (g) Average location survivorship with respect to settlement location at the end of data collection on textured tiles only (n = 6). Throughout, different letters indicate statistically different data.

### Survivorship

Total survivorship at the end of the growth experiment (139 d) was analyzed using an ANOVA, with chemistry and topography as factors. Both chemistry (*P* = 0.03) and topography (*P* < 0.01) affected survivorship (**Figure 3c**). Overall, the 1% and 2% carbonate additions showed the greatest survivorship (**Figure 3c, d**). Across all 15 tiles, the survivorship on flat surfaces was the same across chemistries and topographies (*P* = 0.22), and the survivorship in divots was the same across the 6 textured tiles (*P* = 0.68) (**Figure 3e**). When comparing the survivorship of settlers by location on textured tiles (surface settlers vs. within-divot settlers for each tile), the survivorship of corals settled in divots was significantly lower than the survivorship of corals settled on the flat surfaces of tiles (*P* < 0.01) (**Figure 3f-g**).

### Growth

Surface area, vertical height, and volume of *O. faveolata* varied with respect to chemistry and topography. Additionally, *O. faveolata* surface area and volume were better predicted when including an interactive effect of chemistry and topography in their respective models (**Supplementary Table S5, Table 1, Figure 4**). Both chemistry and topography were found to be significant predictors when generating GLMs of coral growth data with respect to surface area, skeletal height, and volumetric growth. While none of the chemistries tested emerged as the best choice resulting in the greatest growth – each chemistry had its own growth patterns, with topography having a stronger effect on growth than chemistry. *Orbicella faveolata* settlers on the surface of textured tiles have a smaller areal size than on the surface of flat tiles, perhaps due to negative interactions with algae growing in the divots. Indeed, lower growth on textured tiles in general may be because coral growing in divots experience higher competition from other corals and/or algae in these confined spaces or were spatially limited in the area to which they could grow. However, we could not test this directly because our profile imaging for measuring vertical growth could not measure the height of corals growing in divots; individuals settled in divots were excluded from growth analyses.

**Table 1:** GLM Effect tests. Effect tests for significant factors in the best-fit GLM. The best fit GLMs for predicting surface area and volume were dependent on chemistry, topography, and an interactive effect of chemistry and topography, and the best fit GLM for predicting vertical height was dependent only on chemistry and topography (**Supplementary Table S5**).

|  |  | Surface Area | Vertical Height | Volume |
| --- | --- | --- | --- | --- |
| Days in Flume | DF(x,y) | (1, 233) | (1, 249) | (1, 232) |
|  | F | 619.03 | 743.31 | 858.49 |
|  | p | <0.0001 | <0.0001 | <0.0001 |
| Chemistry | DF(x,y) | (2, 231) | (2, 247) | (2, 230) |
|  | F | 5.33 | 8.34 | 3.3 |
|  | p | <0.01 | <0.001 | 0.04 |
| Topography | DF(x,y) | (1, 230) | (1, 246) | (1, 229) |
|  | F | 29.89 | 10.50 | 5.02 |
|  | p | <0.0001 | <0.01 | 0.03 |
| Interactive Effect | DF(x,y) | (1, 229) | NA | (1, 228) |
|  | F | 25.79 | NA | 7.97 |
|  | p | <0.0001 | NA | <0.01 |

**Figure 4:**
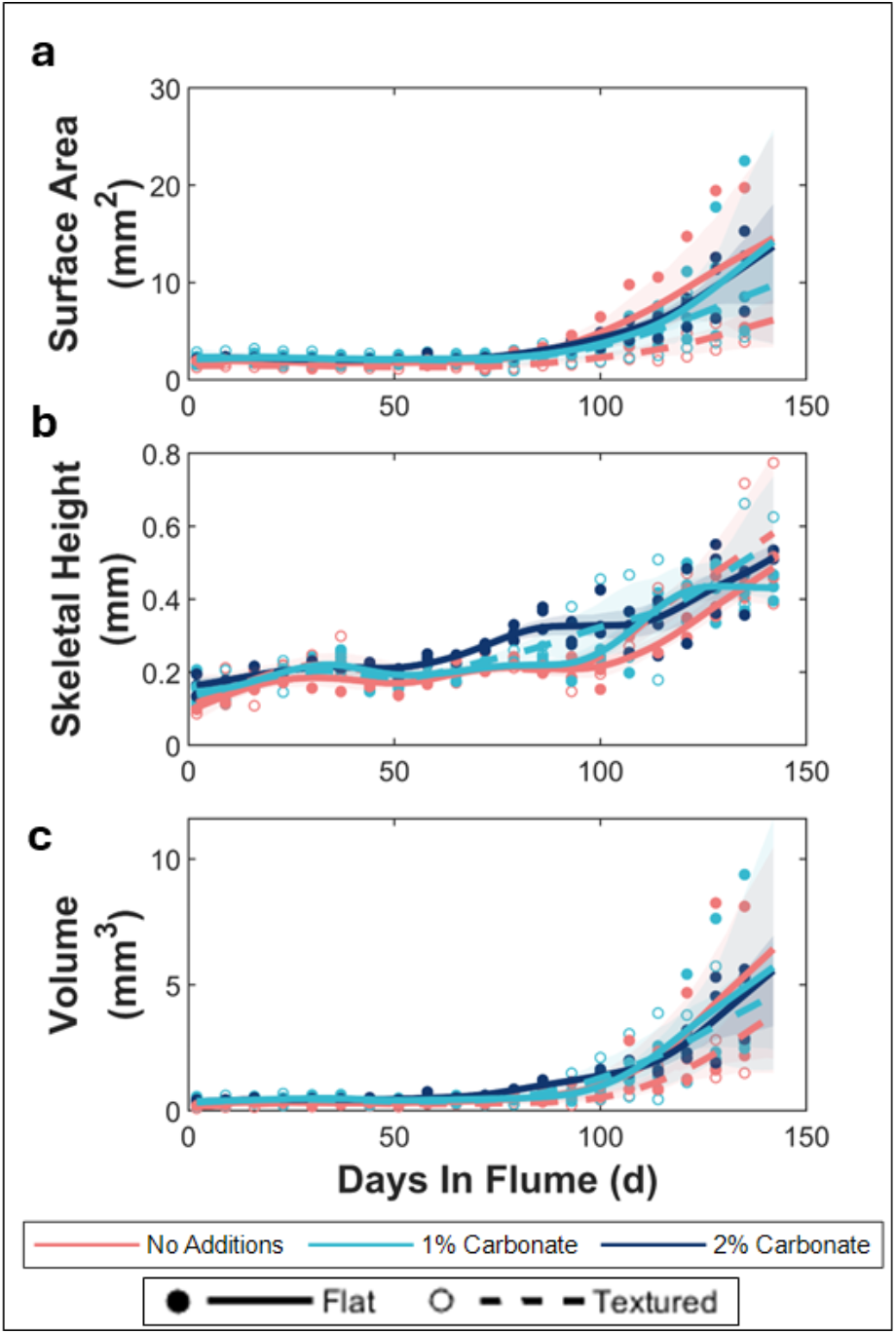
*Orbicella faveolata* growth. Spline-fitted curves and 95% confidence intervals for (a) surface area, (b) skeletal height, and (c) volume of *O. faveolata* settled on the surface of cement tiles. Data has been filtered to only include *O. faveolata* which survived until the end of the data collection period. Chemistry and topography are significant effects in GLMs run on the data.

## Discussion

### Physicochemical Environment of AE Tiles

Adding 1% to 2% sodium carbonate or bicarbonate by weight to cement tiles was sufficient to elevate the pH in the boundary layer by 0.2-0.4 pH units under stagnant flow. When flow was introduced, 1% carbonate and 1% bicarbonate tiles yielded the largest increase in pH (0.1 units) across all chemistries (**Figure 2**). This smaller increase of pH in the advection setting compared to diffusive setting for all chemistries is expected because the carbonate and bicarbonate ions leaching out of the cement will be diluted by the much larger pool of carbonate and bicarbonate ions in the total seawater. The 2% carbonate tiles have a significantly smaller impact on local pH in flow conditions compared to diffusive conditions in both the advection verification and in the aging experiments **(Figure 2)**.

The reduced effect of 2% carbonate addition tiles on pH in the flow compared to diffusion is unexpected. However, the similar ΔpH values for 2% carbonate on the surface and above the surface indicate a more uniform chemical environment above the tile. This becomes apparent when estimating the Péclet number (Pe) for carbonate ions in this environment. Pe represents the ratio of the advective transport rate to the diffusive transport rate of a substance and is calculated as

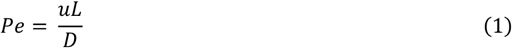

where *u* represents the flow speed (0.012 m s^-1^), *L* is the characteristic length scale (0.03 m, the length of the flume), and *D* is the diffusion coefficient for a given substance, which can be estimated as

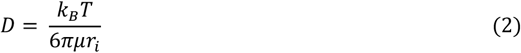

where *k*_*B*_ is the Boltzman constant (1.38 x 10^-23^ J K^-1^), *T* is the absolute temperature of the system (293 K), *μ* is the dynamic viscosity (8.9 x 10^-4^ Pa s), and *r*_*i*_ is the radius of ion in question. When Pe < 1, diffusion dominates transport, when Pe = 1, advection and diffusion are of equal importance, and when Pe > 1, advection dominates transport (16). Here, Pe values for carbonate (*r*_*i*_ = 178 pm) and bicarbonate (*r*_*i*_ = 156 pm) are both much greater than 1 (Pe_Carbonate_ = 2.0 x 10^5^, Pe_Bicarbonate_ = 2.3 x 10^5^), indicating that advection is dominant over diffusion within the boundary layer. This is observed in the weaker effect that tiles have on pH when in advective conditions compared to diffusive conditions.

Advection influencing the tile’s ability to affect pH also can explain why 2% carbonate addition tiles do not perform as well as 1% bicarbonate and 1% carbonate tiles in flow conditions. The change in pH of 2% carbonate tiles in the aging experiment at the surface was not different than the change in pH above the boundary layer. However, 2% carbonate addition tiles significantly raise the pH above ambient conditions. This may be because the 2% carbonate tiles leached their carbonate ions into the environment at a rate where advective mixing no longer stratifies the pH below and above the boundary layer, as it did for the 1% bicarbonate and 1% carbonate addition tiles.

### Coral response to AE Tiles

Initial settlement of *O. faveolata* was not strongly affected by chemistry; however, settlement was affected by topography. *Orbicella faveolata* preferred settling on textured tiles rather than flat tiles, and within textured tiles, a strong preference to settle within the confines of a divot rather than on the surface (**Figure 3**). Coral recruits are more likely to settle in topographical features close to their size to maximize their area for attachment, a phenomenon coined “attachment point theory” (17, 18). In this experiment, the 3 mm diameter of the divot was approximately six times the diameter of recruits, which provides more surface area as the coral settler can attach to the divot bottom or the wall.

Although *O. faveolata* preferred settling on textured tiles, and specifically, within the divots on textured tiles, *O. faveolata* suffered from higher mortality rates on textured tiles, particularly those that settled in divots, than flat tiles, even with AE additions. It is possible the divots were too sheltered from the surrounding environment. Minimal flow was observed in the divots, creating relatively static flow conditions. With no flow to fuel advective processes, Pe << 1, and diffusion dominated the chemical landscape within the divots. A purely diffusive environment limits access to ambient nutrients introduced by the flow and restricts the environment’s ability to flush waste products away from the coral. Although corals generate ciliary flows for nutrient turnover (19), these local flows cannot generate the magnitude of turnover required to flush carbon dioxide and other waste products away while bringing in nutrients and dissolved oxygen for growth. Minimal flow in the divots also allows pH to be significantly higher in the divots compared to the surface. While the intention of the experiment was to expose coral to higher alkalinity conditions, it is possible that the chemical environment within the divots was too extreme. Another consequence of using the flumes to grow coral was that the tiles had to be cleaned by hand rather than co-reared with herbivores to control algal growth. Scraping algae out of the divots during cleaning was a delicate process in such a tight space while not harming coral. Moreover, the algae in the divots could have served as additional competition for resources in an already resource-limited environment (20).

Coral survivorship over the 139-day growth experiment, regardless of settlement location, increased by a factor of 2.5 to 2.9-fold on the 1% and 2% carbonate tiles compared to the tiles not receiving any (**Figure 3**), however, growth had minimal variation with respect to chemistry (**Figure 4**). We initially hypothesized that any increase in survivorship would be a result of increased growth. Ocean acidification makes it difficult for calcifying organisms to build their carbonate skeletons, particularly during early stages of development, by raising the energetic demands for growing skeletons (21-24). For corals, the first deposition of CaCO_3_ for skeletal growth occurs during the first week or two after metamorphosis. The quicker this initial deposition occurs, the greater the success of the organism surviving to reproductive maturity (24).

When planning for these experiments, we envisioned that AE tiles would elevate pH in the laminar boundary layer and corals would benefit from that environment while they were small enough to occupy that zone by growing faster. The observation that survivorship is improved on AE tiles without significantly faster growth suggests that corals benefit from the AE tile chemistry without augmenting their skeletal growth. It is possible that corals survived better because their energy was reallocated to other important processes rather than skeletal growth that were unaccounted for in the experiment, including tissue growth, or it could be that skeletal density rather than areal extension was higher under AE enhancement, and this enhances attachment strength and eventual survivorship.

Corals may also benefit from tile chemistry through direct contact with the substrate, in addition to benefiting via flow. This is evidenced by the enhanced survivorship on tiles that had the least effect on pH under flow – 2% carbonate addition tiles (**Figure 3**). Hence, carbonate and bicarbonate ions may leach from the substrate directly into coral tissues from the point contact under the corals, and the elevation of pH in the boundary layer may be less important.

## Conclusion

In summary, our results show that AE substrates can be an effective method of improving early coral recruit survivorship in laboratory settings. These AE substrates could be potentially attached to artificial reef base structures in the ocean to accelerate the development of coral reef biota on the structure. In addition, settlers grown in land-based spawning and propagation facilities could benefit from higher survivorship on AE tile substrates prior to being outplanted to natural reef surfaces. While field-based demonstrations are needed to demonstrate the scalability of AE substrates for reef restoration and hybrid reef deployments, our results provide promising indications that new technical advancements, such as those presented here, can aid in meeting the coral restoration challenge while also helping build coastal resilience.

## Materials and Methods

### Tile Fabrication (Chemistry)

Three different chemical modifications to Portland limestone cement were tested in addition to unmodified cement with 0.4 water-to-cement ratio. The chemical modifications utilized were 1% sodium bicarbonate, 1% sodium carbonate, or 2% sodium carbonate by weight to cement. Chemical modifications and cement were dry-mixed using a spatula for 4 min to achieve homogeneity, followed by mixing with water for an additional 4 min. All mixtures with added chemicals showed minor reductions in workability from visual observations. Tiles (3 cm x 3 cm x 1 cm) were cast from these four types of AE cement – no additions, 1% bicarbonate, 1% carbonate, and 2% carbonate – referred to as “chemistries”. Two different surface topographies were fabricated using 3D printed female molds – one where the surface of the tile contained no topographical modifications (a “flat” tile), and one where the tile featured a 4 x 4 grid of circular, 3 mm-diameter, 3 mm deep divots spaced 8 mm from each other (divot center to divot center), centered on the tile (a “textured” tile; **Figure S1b**). Molds were filled with paste and placed on a vibration table to remove entrapped air, compacting the mix. The tiles were demolded 24 h after casting by drilling a hole in the bottom of the mold and applying compressed air. Subsequently, tiles were cured in a high relative humidity room (> 95% relative humidity) at 23 °C for 28 d to ensure adequate hydration of cementitious materials.

After curing, tiles were transported from the University of Miami main campus (Coral Gables, FL) to the Rosenstiel School of Marine, Atmospheric, and Earth Science campus (Key Biscayne, FL). Tiles for diffusion experiments were stored in plastic bags until use. Tiles for advection and aging experiments were stored in a 38 L saltwater tank with a bubbler to ensure constant mixing. Tiles for biological experiments were further conditioned for 30 d in running seawater with live rock for colonization by crustose coralline algae (CCA) and biofilms that would provide attractive cues for *O. faveolata* larvae to encourage settlement (25, 26).

To characterize the chemical effect of the four different AE cements, pH was measured using UNISENSE microelectrodes (pH-N; 1.1 mm x 40 mm needle electrode with external reference electrodes, **Figure S2a**). To estimate the diffusive potential of the AE tiles and confirm that they change pH, one randomly selected flat tile of each of the four chemistries was measured continuously over a 24 h period to quantify the diffusive capacity of the tiles to change pH in the water column. For measurements, the tile was transferred to 1000 mL of water. Microelectrodes were positioned at the surface and 2 mm above the surface of the tile. Sensors sampled at a frequency of 1 Hz for 24 hrs.

### Simulating the Hydrodynamic Landscape (Flow + Chemistry)

The hydrodynamic and chemical landscape quantifications in this experiment required the design of a flume that could: (1) achieve flows representative of natural coral reef environments (27, 28), (2) be used in particle image velocimetry experiments to quantify flow fields, and (3) be used in chemistry experiments to easily observe the effects of AE tiles without needing to make changes to the bulk water chemistry. For reasons (2) and (3), we utilize two configurations of a modular flume – a flow-through configuration and a recirculating configuration (15). Both configurations feature the same experimental chamber where measurements are carried out. The flow-through configuration is open-ended, allowing for a constant flow of water through the experimental chamber and the recirculating configuration is fully enclosed, allowing for easy PIV measurements. A brief description of the modular flume, configurations, and flow conditions is provided here, with a more detailed description of the flume’s design, testing, and validation in Ruszczyk, *et al*. (15)

The experimental chamber in both the flow-through and recirculating configurations is comprised of a 30 cm x 10 cm x 14.5 cm transparent acrylic tank with inlet and outlet points made from 3 in, schedule-40 PVC pipe (**Figure 1b**). A 5 cm flow straightener with a hexagonal grid (hexagon circumscribed diameter = 1 cm) is mounted in the upstream direction of the experimental chamber. A 29 cm x 9.8 cm x 1.9 cm plate made of ¾ in PVC was made to hold a 1 x 3 configuration of three tiles along the length of the experimental chamber. Three indentations of dimensions 3.2 cm x 3.2 cm x 1.3 cm were made in the center of the plate, separated by 4.5 cm centered in the middle, to recess tiles such that the surface of the tiles was exposed to flow. The top of the tile plate rested flush with the bottom of the PVC inlet and outlet pipes. In the flow-through and recirculating flume configurations, the experimental chamber was filled such that the surface of the water was flush with the top of the inlet and outlet PVC, resulting in a volume of 2.28 L in the acrylic experimental chamber.

To create a flow-through configuration (**Figure 1c**) where water is constantly turned over, the flumes were connected to a saltwater tap that supplied seawater filtered from the neighboring Biscayne Bay. Four flow-through flumes were installed on one tap line, branching off from a ¾ in PVC mainline. A ball valve at the top of each individual flume’s line allowed for volumetric flow control over each flume placed on the line. After the ball valve, the water flowed through a 1.9-19 L min^-1^ (0.5-5 gal min^-1^) clear in-line flow meter (Omega Engineering) to observe the incoming volumetric flow rate. The water then travelled through a 0.60 m section of 3 in PVC pipe before reaching the inlet of the experimental chamber. The outlet flow was controlled by adjusting a 3 in PVC elbow joint fixed to the outlet of the experimental chamber, and was set such that surface of the water was flush with the top of the inlet and outlet pipes of the experimental chamber.

The recirculating configuration was made by connecting the inlet and outlet pipes of the experimental chamber via a 3 in PVC pipe. An inverted tee joint at the back of the track allowed for a DC motor (Almencla RC Jet Boar Motor Engine Propellor) mounted to PVC pipe to be inserted. The motor was powered by an external power supply to achieve various flow rates. A laminar flow in the experimental chamber with a bulk flow speed of 1.20 cm s^-1^ was generated (Reynolds number = 3550) in all flow-based experiments, matching reef-like conditions (29) (**Figure 1d**).

To estimate the effect of AE tiles in flow conditions, the pH at the tile surface for the four different chemistries was measured in a flow-through flume. One tile of each chemistry was transferred to a flow-through flume and placed in the upstream tile position. Sampling took place via UNISENSE pH microelectrodes positioned at the surface of the tile and approximately 1 cm above the surface of the tile (**Figure S2**). Microelectrodes measured the pH at 1 Hz for 6 h to generate a time series of the influence of chemistries on the local pH at and above the tile surface.

To estimate the longevity of the AE tiles’ enhancement effect, 12 flat cement tiles – three of each chemistry – was measured in an 84-day aging experiment. Tiles were stored in a 3 x 4 grid in a surplus flow-through flume filled with seawater and a bubbler to ensure mixing while experiments were not occurring – a storage flume. The storage flume was connected to the experimental flume such that when experiments occurred, all tiles were exposed to experimental flow conditions. The pH was measured at the surface of the tile and 3.26 ± 0.09 mm above the surface of the tile to capture changes at various heights within the laminar boundary layer. AE tiles were measured in the upstream slot of the tile-holder for 90 min in a flow-through flume with a volumetric flow rate of 0.05 L s^-1^ (1.20 ± 0.3 cm s^-1^) for one of each of the four different chemistries daily for 30 d. After the first 30 d, experiments were performed every other day. The experimental tiles on a given day were randomly selected with no replacements until all 12 tiles had been recorded. After all tiles were measured, the tile position in the storage tank was shuffled, and all tiles were allowed to be selected for experiments again. This ensured a sample size of n = 3 for each of the four different chemistries.

Microelectrodes and reference electrodes were held in the experimental flume via dual micromanipulators (UNISENSE). Microelectrodes were positioned at the center of the downstream edge of the tile (**Figure S2b, c, d**) and the reference electrodes and temperature sensor were placed downstream in the experimental flume to not disturb the flow above the experimental tile. At the beginning of each day, flow was turned on to the flow-through system for the storage flume and the experimental flumes. Microelectrodes were positioned in the tank and allowed to acclimate for up to 30 min or until they provided a steady pH reading of ambient fluid for 10 min. After this acclimation period, a tile was added to the upstream tile position in the tile plate, and microelectrodes were deployed to record data. Microelectrodes ran for 90 min. After this time, the tile was removed, and the microelectrodes were allowed to reacclimate to ambient water for 10 min before the next tile was added. This was repeated such that one tile of each chemistry type was measured per day. Microelectrodes were recalibrated with buffer solutions of standard pH values once per week.

The effect of topography on chemistry was tested by concurrently sampling the pH of textured tiles on the surface of the tile and within a divot on the same tile in a flow-through flume (**Figure S2e**). The first microelectrode was positioned to measure at the bottom of the divot, and the second was positioned within 10 mm of the divot such that it was touching the surface of the tile. Microelectrode positions were measured by photographing the profile and the top-down angle with known reference measurements. One tile of each of the four different chemistries was measured three times over the course of three days for 90 min each day to account for any daily variation in the tile.

The hydrodynamic landscape over flat and textured tiles was quantified in the recirculating flume using 2-dimensional planar particle image velocimetry (PIV). To record the flow over different tile topographies, a tile was placed in the upstream position of a tile-holder. The recirculating flume was seeded with 60 μm diameter polyamide particles (LaVision) and illuminated with a 1350 lumen LED illumination unit (LaVision) mounted above the tank creating a light sheet (**Figure S3**). The thickness of the light sheet was reduced to 1 cm by manually blocking the light entering the tank. The flow was set to a bulk flow velocity of 1.20 cm s ^-1^ using pre-established settings. Once the flow was steady, a 10 s video of the flow above the tile was recorded. Recording took place using an Imager CS2 5 Camera mounted with a L 60 mm focal length lens (F/2.8, 2:1 macro, LaVision) at 40 Hz. Flow quantification over tile topographies took place over an 8 cm x 6 cm window parallel to flow bisecting the tile, resulting in a resolution of approximately 320 px cm^-1^.

### Orbicella faveolata *Settlement and Growth on Tiles (Biology + Flow + Chemistry)*

To test how coral recruits respond to AE cement tiles, *O. faveolata* larvae were settled and grown on AE tiles. Not all chemistries, or combinations of chemistry and topography type could be tested due to spatial limitations in the flumes. We excluded bicarbonate additions for the biology experiment because bicarbonate is not as effective at uptaking H^+^ as carbonate, therefore maximizing the chance of observing any differences in growth related to chemistry. The three chemistry treatments were no additions, 1% carbonate additions, and 2% carbonate additions.

Each chemistry included a set of flat tiles, and the no additions and 1% carbonate additions chemistries included a set of textured tiles, resulting in 5 different treatments of chemistry and topography used in the growth experiment.

*Orbicella faveolata* eggs and sperm were collected at Horseshoe Reef (25.1399 °N, 80.2946 °W) in Key Largo, FL, during a natural spawning event on August 7-8, 2023, and fertilized in-vitro. The larvae were transferred to three polystyrene settlement bins at a density of approximately 300 per bin. Each bin contained a random assortment of 25 treatment tiles, and 5 mL of crushed *Hydrolithon boergesenii* (26) in UV-sterilized, 1 μm filtered seawater. Settlement bins were kept in a temperature-controlled incubation room at 27 °C and daily 75% water changes were conducted while the larvae settled on the cement tiles over 5 weeks. Larvae were characterized as settled upon achieving full metamorphosis on the cement tiles, distinguished by a more circular, squat morphology in contrast to the elongated shape of freeform larvae. *S*ettlers were counted and spatially mapped across all experimental tiles using a dissection microscope (**Figure S4a**). Tiles were labeled using waterproof paper tags affixed on the side of each tile using CorAffix gel coral glue. The three tiles of each of the five treatments with the highest settlement densities on the upper surface were selected for grow out in the flow-through flumes, totaling 180 coral recruits across 15 tiles.

*Orbicella faveolata* were inoculated with symbionts three times per week for 10 weeks. Inoculation occurred 5 weeks prior to relocation in flumes and the first 5 weeks residing in the flumes. *Orbicella faveolata* were inoculated with *Durusdinium* algal culture isolated from Key Largo, FL in the early 2000s. *Durusdinium* cultures were maintained in an incubator at 27 °C in f/2 media with a 14:10 light:dark photocycle under 40 μE light at a density of 10^6^ cells mL^-1^. To inoculate *O. faveolata*, tiles were transferred from their respective tanks to polystyrene bins containing 3,000 cells mL^-1^ of *Durusdinium* algal culture for 6 hrs. Inoculation continued until each polyp on the tiles exhibited color indicative of symbiont saturation. During the inoculation period, tiles received 30 μmol s^1^ m^-2^ of photosynthetically available radiation (PAR) on a 12:12, light:dark photocycle.

Tile position within the flume was randomized 3 times per week to reduce possible downstream effects from the AE tiles. Tile groups were assigned to different flumes twice per week to ensure randomness across flumes. Twice per week, tiles were relocated to 19 L tanks in a 28 °C, temperature-controlled bath for 45 min, and fed with Golden Pearls coral food. In traditional coral tanks, herbivores are co-reared with corals to control algae growth. Herbivores were not used in this experiment due to unsteadiness in flows around the additional organisms, and the threat that they would maneuver outside of the experimental chamber in the flow-through flumes. Instead of relying on herbivores to control algae growth, tiles were manually cleaned twice per week using a 25-gauge needle under a dissection microscope to ensure all algae were removed from the surface of the tiles.

To consistently identify corals throughout the experiment, settlement maps were made by taking an areal image of the full tile and updated weekly. Each recruit was assigned a unique ID, comprised of its tile number and a letter. Settlement maps provided confidence in individual coral measurements for the entirety of the experiment across multiple researchers collecting data.

Coral measurements occurred weekly. Survivorship was visually assessed by observing polyp activity during cleaning and measurement periods. For areal measurements, tiles were transferred to a 100 mL dish and imaged under a dissecting microscope. Images included a ruler in the background for scaling. For vertical growth measurements, corals were transferred to the recirculating flume with no flow present and imaged in a vertical profile using the PIV camera. Corals were located on the tile by mounting a calibration wand to a micromanipulator positioned above the tank. Referencing the settlement maps, the calibration wand was moved to the approximate position of the coral. The image was refined by adjusting the lens on the camera until the coral was in focus. The calibration wand was then physically adjusted to be in focus with the coral, ensuring that a known reference measurement was present in each image. As the image library was generated, old photos were cross-referenced to ensure consistent camera positioning throughout the experiment (**Figure S4c-d**).

*Data processing*

Chemistry data for tiles was processed using a custom written MATLAB code. Chemical data collected from microelectrodes was first transformed from the logarithmic pH measurements to linear [H^+^] measurements by

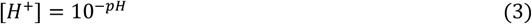

allowing for processing and analysis using linear techniques. To account for slight differences in calibration between microelectrodes and daily differences in ambient pH in experiments, the change in [H^+^] (*ΔH*^*+*^) is used, rather than absolute [H^+^] measurements. To calculate *ΔH*^*+*^, the time series data was transformed such that the average [H^+^] in the 10 min calibration period of each time series is equal to zero. To account for sensor drift in the microelectrodes, data was detrended using a natural logarithmic function fitted to the calibration period of the data.

Flow data for tiles was processed in DaVis 11.0.0.196 software (LaVision). For PIV over tile topographies, a mask was applied to the bottom of the tank to exclude any areas where the tile may have protruded above the tile-holder. PIV was performed using a bicubic interpolation to interpolate vectors, 5 x 5 denoising, a symmetrical shift correction mode. Pixels were interpolated using a spline interpolation mode with a window interrogation size of 16 px x 16 px, a direct correlation algorithm was used between frames, and cell size weighted as a round cell. These parameters resulted in an average correlation value of 1.00 px.

*Orbicella faveolata* data was first scanned to exclude any chimeras (n=5) – instances of two adjacent settlers fusing to form one large entity – from the analysis. Data was analyzed using three response variables: settlement preference, survivorship, and growth. Settlement preference was assessed by counting the total number of *O. faveolata* settlers per tile, parsed by settlement location (i.e., on the flat surface or within a divot). Two different measures of survivorship were calculated and analyzed. Total survivorship with respect to the tile was calculated each week as the ratio of all surviving *O. faveolata* on a given tile to the initial number of settlers on that tile, regardless of settlement location. Survivorship was partitioned for textured tiles into the ratio of surviving *O. faveolata* settlers on flat surface versus within divots, relative to the initial number of settlers in those locations – location survivorship. For flat tiles, only survivorship of flat surface settlers was calculated, as there were no divots present.

Polyp surface area was calculated by analyzing images in ImageJ software (version 1.54g). The polyps in photos were outlined twice with the freehand tool and the area was calculated and measured twice. The radius of the coral, *r*, was estimated from this area, assuming the fitted shape was a circle. Vertical measurements were analyzed in DaVis 11.0.0.196 software. Vertical height was measured from the surface of the tile to the visible skeletal height, identifiable by a change in contrast in the image. Volumetric growth was calculated using the areal and profile images. Coral volume (mm^3^) was modelled as a cylinder, using the radius of the fitted circle calculated in areal measurements and the skeletal height of the coral as

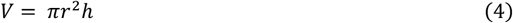

where *r* is the radius (mm) of the areal measurements and *h* is the skeletal height (mm) of the coral (**Figure 1b**).

### Statistical Analysis

Statistical analysis for chemical characterization of tiles was performed on *ΔH*^*+*^ to allow for analysis using linear techniques. Results reporting chemical differences are presented as *ΔpH* values, for ease of understanding. The influence of tiles on local pH via diffusion and advection was assessed using an ANOVA on one 6 h time series for each chemistry type for each flow condition. Aging data was modelled using linear regressions fitted to the average daily *ΔH*^*+*^ at the surface and above the surface for each chemistry. The effect of topography on chemistry was assessed using an ANOVA on one 1.5 h time series for each chemistry.

Flows over surface topographies were compared via velocity and vorticity measurements at similar spatial coordinates. The data was checked for outliers resulting from PIV analysis, and outliers were removed from the dataset.

*Orbicella faveolata* data was analyzed with respect to tile. Settlement preference was analyzed by the initial number of settlers (0 d) using a factorial ANOVA, with chemistry and topography as factors. *Orbicella faveolata* total survivorship was analyzed at the conclusion of the experiment (139 d) using a factorial ANOVA with chemistry and topography as factors. On the textured tiles, where larvae had the option of settling in different settlement locations (i.e., on the surface or within a divot), the percentage of settlers based on location (survivorship with respect to location) for the textured tiles was calculated as the amount settled in each location per tile (i.e., on the exposed surface or in a divot) and was analyzed via Student’s t-test with settlement location as a factor.

To remove any selective bias in growth data from smaller corals dying, growth data was filtered to only include colonies which survived the duration of the experiment. Growth analysis only included measurements for corals grown on the flat surfaces of tiles, as vertical measurements could not be obtained for corals which settled in the divots of textured tiles. Growth data (surface area, skeletal height, and volume) was analyzed using generalized linear models (GLMs) (30). GLMs analyzed growth data with respect to age (days in flume) and a factorial design of chemistry and topography using an inverse function linked to a gamma distribution. AIC values of each model were compared within dependent variables to determine the best fitted model (30).

Statistical analysis was performed in JMP 16.0 (SAS Institute 2023) and R 4.4.1 (R Core Team 2022). Assumptions for normality and heteroscedasticity were verified on all the data, allowing for parametric ANOVAs and Tukey’s HSD using the DHARMa package (31) in R. All data is presented as mean ± standard error unless otherwise stated.

## Acknowledgments

The authors would like to acknowledge all members of the X-REEFs team (Reef Engineering to Enhance Future Structures project, funded by the Defence Advanced Research Projects Agency) for their support. Additionally, the authors thank Dr. Sanchit Mehta and the students in the SUSTAIN laboratory at the University of Miami, Johnnia Xia, Owen Brown and Patrick M. Kiel in the Prakash Lab and Cedric M. Guigand (Rosenstiel School Maker Shop) for flume development, design, and initial testing, Clara Haughey-Gramazio in the Langdon Lab for coral assistance, Dr. Mary Alice Coffroth for *Durusdinium* cultures (University at Buffalo), as well as all members of the Suraneni, Baker, Langdon, and Prakash Labs for useful discussions. The authors also thank Dr. Douglas Neal from LaVision Inc, for installation and support of the PIV system. V.N.P. would like to acknowledge start-up funding support from the University of Miami.

This material is based upon work supported by the Defense Advanced Research Projects Agency under the Reefense Program, BAA HR001121S0012. The views, opinions and/or findings expressed are those of the author and should not be interpreted as representing the official views or policies of the Department of Defense or the U.S. Government.

