## Supporting Information File for "Local alkalinity enhancement using artificial substrates increases survivorship of early-stage coral recruits"

**This PDF file includes:**

Figures S1 to S5
Tables S1 to S5

**Supplementary Figures/Tables:**

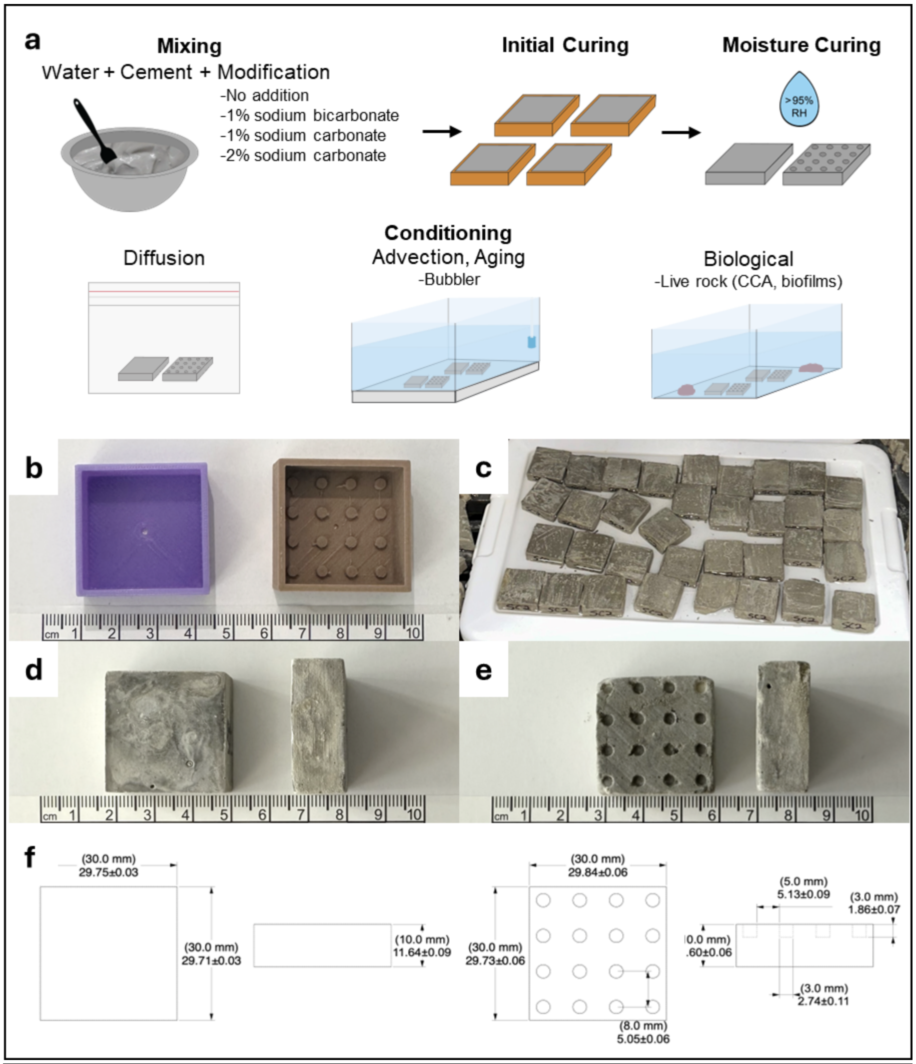


**Figure S1: AE Tile fabrication methods.** (a) Cartoon of AE tile fabrication process. To make AE tiles, dry mix containing cement and modifications was added to water and mixed for 4 min. Paste was cast into 3D-printed molds and cured for 24 h in ambient conditions for its initial cure. Tiles were then demolded and cured inside a humidity room (23 C and >95% relative humidity) for an additional 27 days. Final conditioning for AE tiles varied based on intended experiment. Diffusion tiles were stored in plastic bags. Advection and aging tiles were stored in a tank with a bubbler. Tiles for biological experiments were conditioned in running seawater and live rocks for colonization. (b) 3D-printed female molds with two distinct surface topographies – flat and textured. (c) Tiles curing in >95% humidity room. (d) Top and side view of sample flat tile. (e) Top and side view of sample textured tile. (f) Tile dimensions for each topography type including designed measurements and actual measurements ± standard error (n = 4).

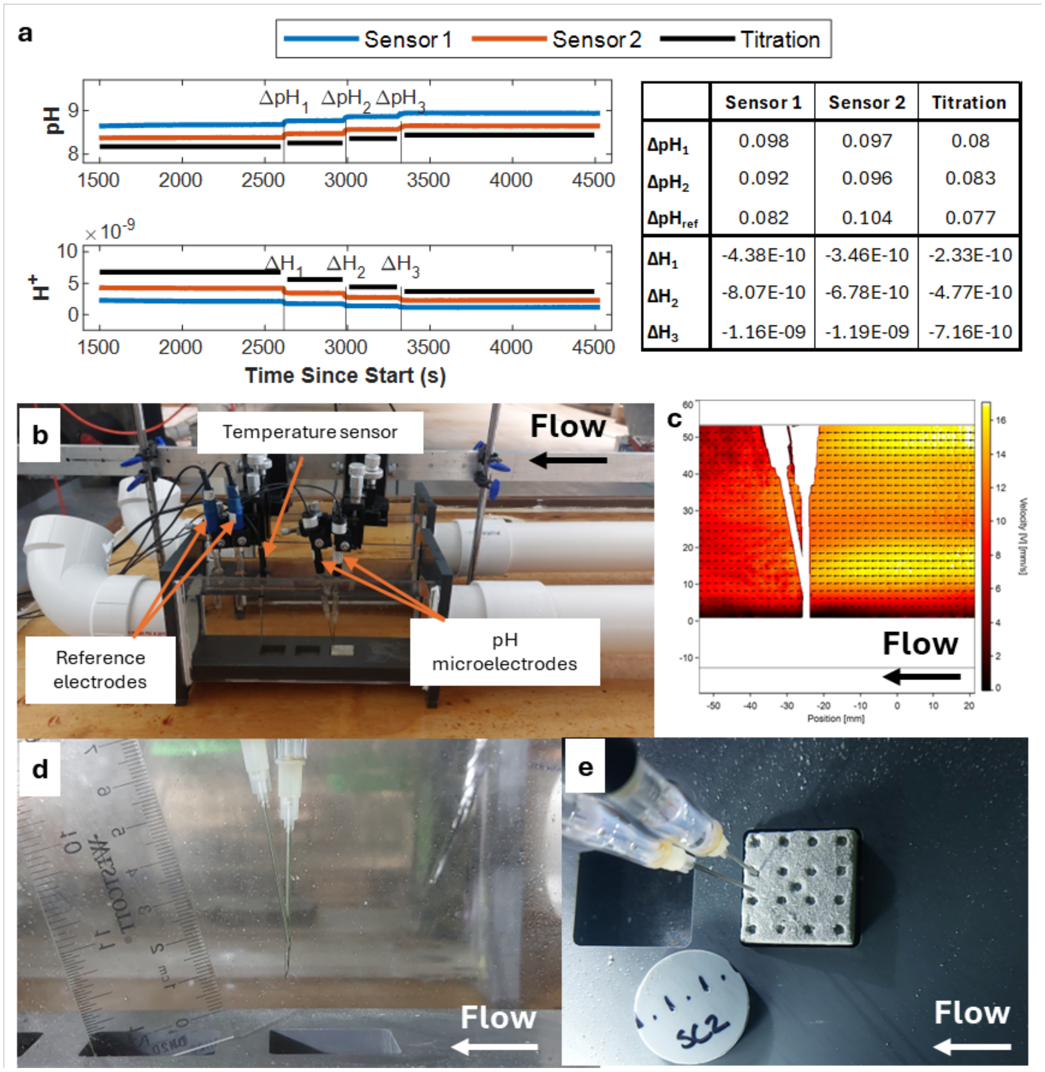


| | Sensor 1 | Sensor 2 | Titration |
|---|---|---|---|
| $\Delta pH_1$ | 0.098 | 0.097 | 0.08 |
| $\Delta pH_2$ | 0.092 | 0.096 | 0.083 |
| $\Delta pH_{ref}$ | 0.082 | 0.104 | 0.077 |
| $\Delta H_1$ | -4.38E-10 | -3.46E-10 | -2.33E-10 |
| $\Delta H_2$ | -8.07E-10 | -6.78E-10 | -4.77E-10 |
| $\Delta H_3$ | -1.16E-09 | -1.19E-09 | -7.16E-10 |

**Figure S2: Chemical quantification of tile measurements.** (a) Sensor accuracy was compared to the standard titration method of pH quantification. Sensors measured 1L of saltwater on a magnetic stir plate to ensure a well-mixed environment. Aqueous bicarbonate was added at time points indicated by vertical lines in the figure on the right and the pH of the solution was measured by both microelectrodes and a sample was removed for measurement by titration. Each method reported slightly different pH and $H^+$ values, however, the change in pH and H+ between time points was similarly low across all methods of measuring pH (table, right). (b) Standard placement of sensors for chemical measurements in flow through flumes. (c) Velocity field of flow around sensors in flow conditions. The flow upstream of the sensors is constant with no mixing. Downstream, minor fluctuations in flow occur due to the sensors. Microelectrodes are angled to sample upstream measurements. (d) Standard sensor position for chemical quantification of tiles in flow conditions for advective quantification and aging experiments. Note the open end of the needle microelectrodes point upstream. (e) Standard vertical sensor position for chemical quantification of textured tiles in flow conditions. Note the one sensor is fully deployed into a divot, while the other sensor is positioned at a similar length along the tile at the surface.

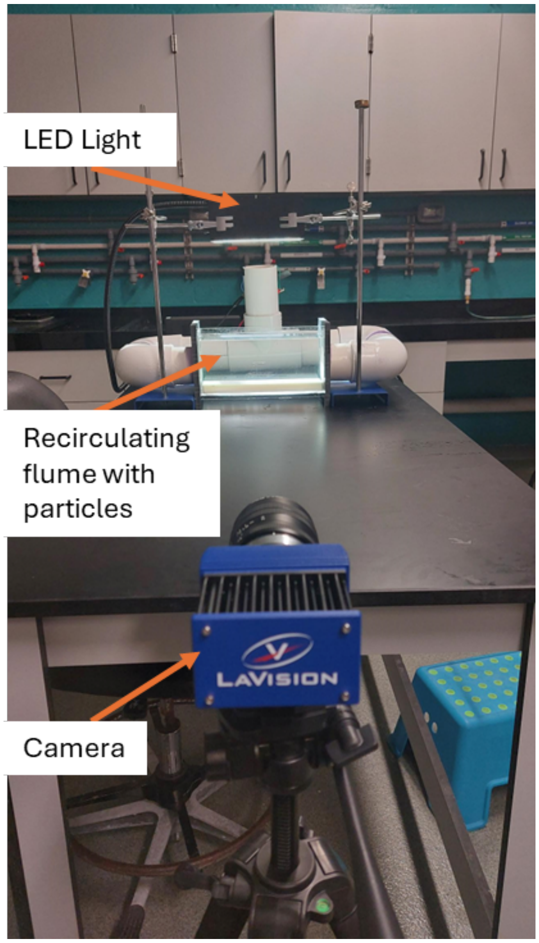


**Figure S3: Particle image velocimetry (PIV) measurements.** Two-dimensional particle image velocimetry (PIV) measurements were taken in a recirculating flume illuminated from above with an LED light source. The flume is seeded with neutrally buoyant particles, and flow is controlled via a propellor at the back of the flume. Videos are recorded with a camera and processed in DaVis (not pictured).

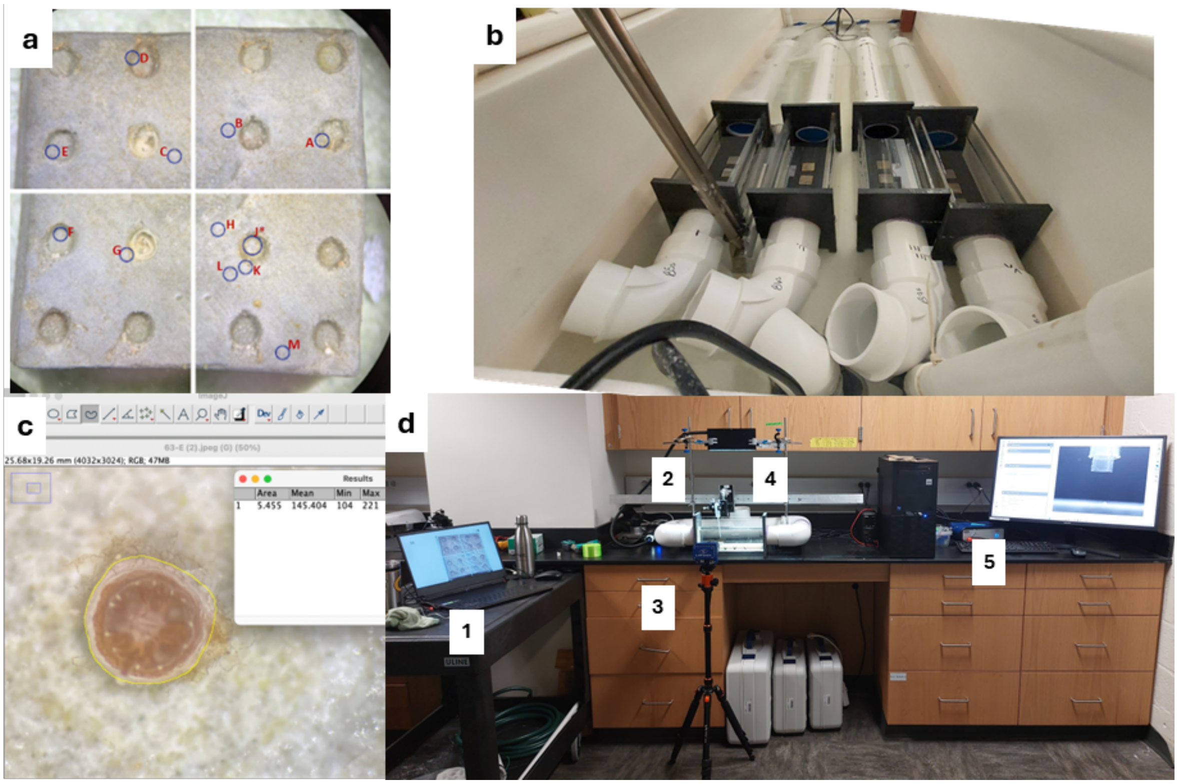


**Figure S4: Coral rearing and measurements.** (a) Four flow through flumes for *O. faveolata* rearing in a temperature-controlled water bath. (b) Sample coral map of *O. faveolata* settlement locations on a textured tile. (c) Lab configuration of vertical growth measurements and steps for measurement. Coral on an experimental tile placed in the recirculating flume can be vertically measured by (1) locating the approximate settlement location of the settler map, (2) moving the calibration wand over the tile such that it matches the approximate location on the settlement map, (3) refining the measurement of the coral by focusing the camera on the coral, (4) finalizing the position of the calibration wand using the micromanipulator such that it is also in focus with the coral, and (5) capturing the image of the in-focus coral and calibration wand in DaVis. (d) Sample areal measurement of *O. faveolata* in ImageJ. Area is calculated using the freehand tool to draw around the perimeter of the coral (yellow).

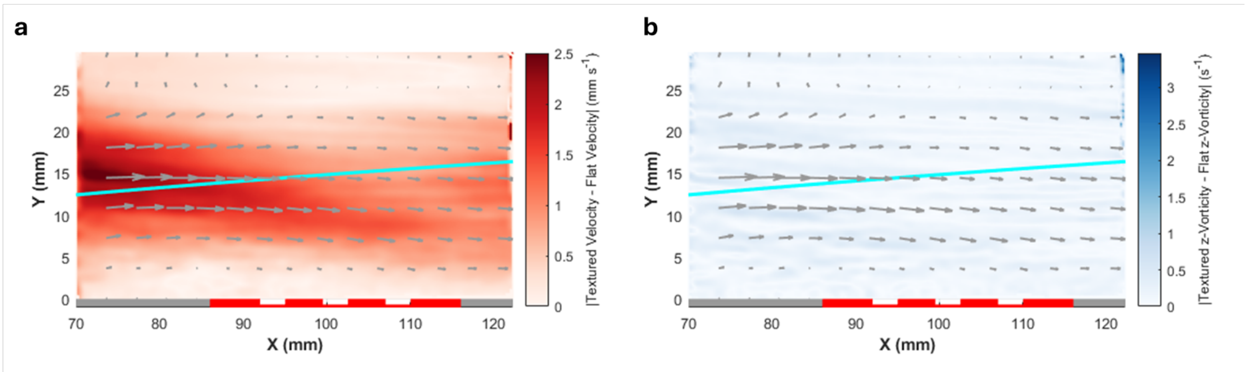


**Figure S5: Topography hydrodynamic comparison.** (a) Difference in velocity between flow over a flat tile and a textured tile. (b) Difference in vorticity between flow over a flat tile and a textured tile. The cyan line on both figures represents the boundary layer calculated from theory.

**Table S1: Chemical Characterization Statistics.** Tukey HSD multiple comparisons for chemical characterization of alkalinity enhanced tiles. Statistics were run on delta $H^+$ values. Adjusted DF = 14403. All 28 unique combinations of chemistry and flow type are tested. PLC = no additions, SB1 = 1% bicarbonate additions, SC1 =1% carbonate additions, and SC2 = 2% carbonate additions. Significant p-values less than 0.05 are bolded in red.

| Comparison | | t | p |
|---|---|---|---|
| Advection x Advection | PLC Advection x SB1 Advection | 9.61E+01 | **<.0001** |
| | PLC Advection x SC1 Advection | 1.58E+02 | **<.0001** |
| | PLC Advection x SC2 Advection | 7.63E+01 | **<.0001** |
| | SB1 Advection x SC1 Advection | 6.18E+01 | **<.0001** |
| | SB1 Advection x SC2 Advection | -1.99E+01 | **<.0001** |
| | SC1 Advection x SC2 Advection | -8.16E+01 | **<.0001** |
| Diffusion x Diffusion | PLC Diffusion x SB1 Diffusion | -3.61E+01 | **<.0001** |
| | PLC Diffusion x SC1 Diffusion | 2.69E+01 | **<.0001** |
| | PLC Diffusion x SC2 Diffusion | 1.01E+02 | **<.0001** |
| | SB1 Diffusion x SC1 Diffusion | 6.30E+01 | **<.0001** |
| | SB1 Diffusion x SC2 Diffusion | 1.37E+02 | **<.0001** |
| | SC1 Diffusion x SC2 Diffusion | 7.44E+01 | **<.0001** |
| Advection x Diffusion | PLC Advection x PLC Diffusion | 1.69E+02 | **<.0001** |
| | PLC Advection x SB1 Diffusion | 1.33E+02 | **<.0001** |
| | PLC Advection x SC1 Diffusion | 1.96E+02 | **<.0001** |
| | PLC Advection x SC2 Diffusion | 2.71E+02 | **<.0001** |
| | SB1 Advection x PLC Diffusion | 7.33E+01 | **<.0001** |
| | SB1 Advection x SB1 Diffusion | 3.73E+01 | **<.0001** |
| | SB1 Advection x SC1 Diffusion | 1.00E+02 | **<.0001** |
| | SB1 Advection x SC2 Diffusion | 1.75E+02 | **<.0001** |
| | SC1 Advection x PLC Diffusion | 1.15E+01 | **<.0001** |
| | SC1 Advection x SB1 Diffusion | -2.46E+01 | **<.0001** |
| | SC1 Advection x SC1 Diffusion | 3.84E+01 | **<.0001** |
| | SC1 Advection x SC2 Diffusion | 1.13E+02 | **<.0001** |
| | SC2 Advection x PLC Diffusion | 9.32E+01 | **<.0001** |
| | SC2 Advection x SB1 Diffusion | 5.72E+01 | **<.0001** |
| | SC2 Advection x SC1 Diffusion | 1.20E+02 | **<.0001** |
| | SC2 Advection x SC2 Diffusion | 1.94E+02 | **<.0001** |

**Table S2: Aging Results Regressions.** Coefficients *a* and *b* for the fitted linear regressions "Delta $H^+$ = a*Day+b" for each combination of chemistry and sensor position. PLC = no additions, SB1 = 1% bicarbonate additions, SC1 =1% carbonate additions, and SC2 = 2% carbonate additions. Significant p-values less than 0.05 are bolded in red.

| | | Fitted Function | | $R^2$ | DF | F | p |
|---|---|---|---|---|---|---|---|
| | | a* | b* | | | | |
| **PLC** | **Above** | 2.629E-02 | -1.017 | 0.12 | (1,52) | 7.17 | **0.01** |
| | **Surface** | 2.293E-02 | -1.153 | 0.05 | (1,52) | 2.85 | 0.10 |
| **SB1** | **Above** | 1.501E-02 | -1.642 | 0.03 | (1,52) | 1.71 | 0.20 |
| | **Surface** | 6.511E-02 | -7.143 | 0.05 | (1,52) | 2.92 | 0.09 |
| **SC1** | **Above** | 2.640E-02 | -1.784 | 0.06 | (1,52) | 3.08 | 0.09 |
| | **Surface** | 5.750E-02 | -8.431 | 0.04 | (1,52) | 2.03 | 0.16 |
| **SC2** | **Above** | 2.415E-02 | -1.616 | 0.08 | (1,51) | 4.73 | **0.03** |
| | **Surface** | 4.455E-02 | -2.802 | 0.13 | (1,51) | 7.54 | **<0.01** |

*Divided by e-10

**Table S3: Aging Stats Comparisons.** Tukey HSD multiple comparisons for aging data. Statistics are run on delta $H^+$ values.  Adjusted DF for pairwise comparisons = 15836, DF for comparisons to 0 = 53. PLC = no additions, SB1 = 1% bicarbonate additions, SC1 =1% carbonate additions, and SC2 = 2% carbonate additions. “Above” refers to above the tile in the bulk flow, “Surface” refers to at the tile surface. Significant p-values < 0.05 are bolded in red.

| Comparison | | t | p |
|---|---|---|---|
| Above x Above | PLC1 Above x SB1 Above | 1.11 | 0.9541 |
| | PLC1 Above x SC1 Above | 0.84 | 0.9909 |
| | PLC1 Above x SC2 Above | 0.74 | 0.9958 |
| | SB1 Above x SC1 Above | -0.28 | 1 |
| | SB1 Above x SC2 Above | -0.37 | 1 |
| | SC1 Above x SC2 Above | -0.1 | 1 |
| Surface x Surface | PLC1 Surface x SB1 Surface | 4.97 | **<.0001** |
| | PLC1 Surface x SC1 Surface | 6.67 | **<.0001** |
| | PLC1 Surface x SC2 Surface | 1.89 | 0.5598 |
| | SB1 Surface x SC1 Surface | 1.7 | 0.6873 |
| | SB1 Surface x SC2 Surface | -3.08 | **0.045** |
| | SC1 Surface x SC2 Surface | -4.78 | **<.0001** |
| Above x Surface | PLC1 Above x PLC1 Surface | 0.28 | 1 |
| | PLC1 Above x SB1 Surface | 5.25 | **<.0001** |
| | PLC1 Above x SC1 Surface | 6.95 | **<.0001** |
| | PLC1 Above x SC2 Surface | 2.16 | 0.3753 |
| | SB1 Above x SB1 Surface | 4.14 | **0.0011** |
| | SB1 Above x PLC Surface | 0.84 | 0.991 |
| | SB1 Above x SC1 Surface | 5.84 | **<.0001** |
| | SB1 Above x SC2 Surface | 1.05 | 0.9657 |
| | SC1 Above x SC1 Surface | 6.11 | **<.0001** |
| | SC1 Above x PLC1 Surface | 0.56 | 0.9993 |
| | SC1 Above x SB1 Surface | -4.41 | **0.0003** |
| | SC1 Above x SC2 Surface | 1.33 | 0.8876 |
| | SC2 Above x SC2 Surface | 1.43 | 0.8442 |
| | SC2 Above x PLC1 Surface | 0.46 | 0.9998 |
| | SC2 Above x SB1 Surface | -4.51 | 0.0002 |
| | SC2 Above x SC1 Surface | -6.21 | **<.0001** |
| Compare to estimated mean = 0 | PLC Above x 0 | -0.43 | 0.67 |
| | SB1 Above x 0 | -3.92 | **<.001** |
| | SC1 Above x 0 | -2.30 | **0.03** |
| | SC2 Above x 0 | -2.75 | **<.01** |
| | PLC Surface x 0 | -1.06 | 0.29 |
| | SB1 Surface x 0 | -5.09 | **<.001** |
| | SC1 Surface x 0 | -6.40 | **<.001** |
| | SC2 Surface x 0 | -2.21 | **0.03** |

**Table S4: Chem X Topography comparisons.** Tukey HSD multiple comparisons for topography x flow data. Statistics are run on delta $H^+$ values. Adjusted DF for pairwise comparisons = 43320. All 28 unique combinations of chemistry and sensor position are tested. PLC = no additions, SB1 = 1% bicarbonate additions, SC1 =1% carbonate additions, and SC2 = 2% carbonate additions. "Surface" refers to the surface of the tile, and "Divot" refers to at the bottom of the divot. Significant p-values less than 0.05 are bolded in red.

| Comparison | | t | p |
|---|---|---|---|
| Surface x Surface | PLC1 Surface x SB1 Surface | 51.78 | **<.0001** |
| | PLC1 Surface x SC1 Surface | 72.7 | **<.0001** |
| | PLC1 Surface x SC2 Surface | -137.1 | **<.0001** |
| | SB1 Surface x SC1 Surface | 21.14 | **<.0001** |
| | SB1 Surface x SC2 Surface | -188.6 | **<.0001** |
| | SC1 Surface x SC2 Surface | -208.8 | **<.0001** |
| Divot x Divot | PLC1 Divot x SB1 Divot | 6.25 | **<.0001** |
| | PLC1 Divot x SC1 Divot | 29.23 | **<.0001** |
| | PLC1 Divot x SC2 Divot | 11.64 | **<.0001** |
| | SB1 Divot x SC1 Divot | 22.98 | **<.0001** |
| | SB1 Divot x SC2 Divot | 5.38 | **<.0001** |
| | SC1 Divot x SC2 Divot | -17.61 | **<.0001** |
| Surface x Divot | PLC1 Surface x PLC1 Divot | -635.6 | **<.0001** |
| | PLC1 Surface x SB1 Divot | -628.4 | **<.0001** |
| | PLC1 Surface x SC1 Divot | -602.3 | **<.0001** |
| | PLC1 Surface x SC2 Divot | -622.5 | **<.0001** |
| | SB1 Surface x PLC1 Divot | 686.4 | **<.0001** |
| | SB1 Surface x SB1 Divot | -679.1 | **<.0001** |
| | SB1 Surface x SC1 Divot | -652.9 | **<.0001** |
| | SB1 Surface x SC2 Divot | -673.2 | **<.0001** |
| | SC1 Surface x PLC Divot | 704.27 | **<.0001** |
| | SC1 Surface x SB1 Divot | 696.98 | **<.0001** |
| | SC1 Surface x SC1 Divot | -670.8 | **<.0001** |
| | SC1 Surface x SC2 Divot | -691.1 | **<.0001** |
| | SC2 Surface x PLC Divot | 497.05 | **<.0001** |
| | SC2 Surface x SB1 Divot | 490.05 | **<.0001** |
| | SC2 Surface x SC1 Divot | 464.74 | **<.0001** |
| | SC2 Surface x SC2 Divot | -484.3 | **<.0001** |

**Table S5: GLM AIC comparison.** AIC values for GLMs run to predict significant factors for growth data. Dependent variables tested include surface area, height, and volume of *O. faveolata*, and independent variables tested include chemistry, topography, chemistry and topography, and chemistry and topography and an interactive effect. The best model (lowest AIC) is bolded.

| | Surface Area | Vertical Height | Volume |
|---|---|---|---|
| **Chemistry** | 711.42 | -764.28 | 125.91 |
| **Topography** | 685.19 | -752.04 | 122.52 |
| **Chemistry + Topography** | 686.32 | **-772.64** | 122.54 |
| **Chemistry + Topography + Interactive Effect** | **662.25** | -772.35 | **115.78** |
